# Inferring *Mycobacterium bovis* transmission between cattle and badgers using isolates from the Randomised Badger Culling Trial

**DOI:** 10.1101/2021.05.27.445931

**Authors:** Andries J. van Tonder, Mark Thornton, Andrew J.K. Conlan, Keith A. Jolley, Lee Goolding, Andrew P. Mitchell, James Dale, Eleftheria Palkopoulou, Philip J. Hogarth, R. Glyn Hewinson, James L.N. Wood, Julian Parkhill

## Abstract

*Mycobacterium bovis* (*M. bovis)* is a causative agent of bovine tuberculosis, a significant source of morbidity and mortality in the global cattle industry. The Randomised Badger Culling Trial was a field experiment carried out between 1998 and 2005 in the South West of England. As part of this trial, *M. bovis* isolates were collected from contemporaneous and overlapping populations of badgers and cattle within ten defined trial areas. We combined whole genome sequences from 1,442 isolates with location and cattle movement data, identifying transmission clusters and inferred rates and routes of transmission of *M. bovis*. Most trial areas contained a single transmission cluster that had been established shortly before sampling, often contemporaneous with the expansion of bovine tuberculosis in the 1980s. The estimated rate of transmission from badger to cattle was approximately two times higher than from cattle to badger, and the rate of within-species transmission considerably exceeded these for both species. We identified long distance transmission events linked to cattle movement, recurrence of herd breakdown by infection within the same transmission clusters and superspreader events driven by cattle but not badgers. Overall, our data suggests that the transmission clusters in different parts of South West England that are still evident today were established by long-distance seeding events involving cattle movement, not by recrudescence from a long-established wildlife reservoir. Clusters are maintained primarily by within-species transmission, with less frequent spill-over both from badger to cattle and cattle to badger.

## Introduction

*Mycobacterium bovis* (*M. bovis*), a member of the *Mycobacterium tuberculosis* complex (MTBC) and a pathogen with zoonotic potential [1], is the main causative agent of bovine tuberculosis (bTB), a significant source of morbidity and mortality in the global cattle industry. In the United Kingdom (UK), the estimated annual cost of managing this disease is £120 million [2].

*M. bovis* has a broad host range with different wildlife reservoirs depending on geographic location: in Britain and Ireland the Eurasian badger is the predominant wildlife host, in France wild boar and deer, and in New Zealand the introduced brush-tail possum [3–5]. The presence of wildlife reservoirs makes the control and potential elimination of bTB challenging even in countries such as the UK with extensive cattle test and slaughter strategies, and movement restrictions imposed on herds with new bTB incidents termed breakdowns [6].

The Randomised Badger Culling Trial (RBCT) was a large-scale ecological field experiment carried out between 1998 and 2005 with the aim of quantifying the impact of culling badgers on the incidence of bTB breakdowns in nearby cattle herds [7]. Ten trial areas within the southwest of England and English Midlands, each of approximately 100 km^2^, were selected on the basis of high bTB incidence. Each trial area was divided into triplets of randomly allocated interventions: proactive culling (widespread and repeated culling across the trial areas), reactive culling (badgers culled if breakdowns detected in nearby herds) and control or survey-only areas (no badger culling). Approximately 9,000 badgers were culled and sampled in proactive areas between 1998 and 2005 though culling was suspended between May 2001 and January 2002 due to a national foot and mouth disease epidemic.

A number of previous studies have established epidemiological links between badgers and nearby cattle although extent of transmission between the two host species remains uncertain [8, 9]. More recent analyses making use of whole genome sequencing (WGS), which offers much higher resolution for strain characterisation and tracking transmission, have confirmed the close genetic relatedness of *M. bovis* isolates from sympatric cattle and badger populations but, due to the low genomic variability of the *M. bovis* genome and a lack of balanced sampling between the different host species, have not been able to adequately address the direction of transmission [10, 11]. The first direct estimate of the extent and directionality of transmission between cattle and badgers suggested that transmission was up to ten times higher from badgers to cattle than *vice versa* [9]. Subsequent studies have estimated that cattle to badger transmission was at least three times or an order of magnitude higher than badger to cattle transmission [12, 13]. However, these results may not be applicable to the wider *M. bovis* population in different regions of the UK, often being based on small, geographically narrow datasets chosen for the presence of the same strain type (spoligotype SB0263). Those in Northern Ireland may additionally reflect the lower density of badgers compared to Southern England and the outbreak in Cumbria was an outbreak in a region with low incidence of bTB.

This Eradication of bovine tuberculosis (ERADbTB) project was set up with the aim of using WGS data obtained from *M. bovis* isolates collected as part of the RBCT to characterise the population structure of the bacterium within the trial areas, attempt to quantify levels and directionality of *M. bovis* transmission between cattle and badgers and track the longer-term persistence of genetic lineages of the bacterium. Approximately 2,000 *M. bovis* isolates available from the RBCT were selected for sequencing with the final dataset consisting of 1,442 genomes (690 from badgers and 750 from cattle found to be infected in proactive cull trial areas respectively).

## Methods

### Sample selection, culturing and sequencing

A total of 2,137 *M. bovis* isolates from cattle (n = 1,011) and badgers (n = 1,126) collected from proactive trial areas were selected for culturing, of which 1,838 isolates were located in the frozen archives maintained by the Animal and Plant Health Agency (APHA). Isolates were re-cultured and grown for up to six weeks or until sufficient growth was observed (n = 1,651). Isolates were heat killed in hot blocks at 80°C for 30 minutes. An adapted library construction protocol using an increased number of sixteen PCR cycles was used to generate Illumina libraries which were then sequenced at the Wellcome Sanger Institute using the Illumina HiSeq X10 platform to generate 2 x 150 bp paired-end reads. Metadata for the sequenced isolates is available on pubMLST (https://pubmlst.org/projects/mbovis-eradbtb) [14, 15]. A map of the geographical locations of isolate collection (latitude and longitude) was constructed using the R v 3.5.1 [16] library ggmap [17].

### Sequence QC

FastQC v0.11.9 [18] was used to generate basic quality control metrics for the raw sequence data. Sequencing reads were prefiltered using Kraken v0.10.6 [19] against a database containing all RefSeq bacterial and archeal nucleotide sequences to identify reads with similarity to *Mycobacterium* species. Further sequence matching was done on the Kraken results using Bracken v1.0 [20]. Samples with < 70% reads mapping to a *Mycobacterium* species were excluded from further analyses (n = 183).

### In silico genotyping

SpoTyping v2.0 [21] was used to extract the binary representation of spoligotype patterns from the sequence reads and the *M. bovis* spoligotype database (https://www.mbovis.org/database.php) was used to assign SB numbers. Novel spoligotype patterns were submitted to the database to generate new SB numbers. Clonal complexes were assigned to samples using RD-analyzer v1.0 [22] with samples not identified as belonging to previously described clonal complexes (Eu1, Eu2, Af1, Af2) designated as “Other” [23–26]. Further assignment of isolates marked as “Other” to clonal complex was based on the phylogenetic lineages recently identified by Loiseau *et al*. [27].

### Mapping and phylogenetics

Sequence reads were mapped to the *Mycobacterium bovis* AF2122/97 reference genome (NC0002945) using BWA mem v0.7.17 (minimum and maximum insert sizes of 50 and 1000 respectively) [28]. Single nucleotide polymorphisms (SNPs) were called using SAMtools v1.2 mpileup and BCFtools v1.2 (minimum base call quality of 50 and minimum root squared mapping quality of 30) as previously described [29]. Samples with reads mapping to less than 90% of the AF2122/97 reference were excluded (n = 26). Genomic regions consisting of GC-rich sequences such as PPE proteins and repeats were masked in the resulting alignment using previously published coordinates [30] and variant sites in the subsequent masked alignment were extracted using snp-sites v 2.5.1 [31]. Maximum likelihood phylogenetic trees were constructed using IQ-tree v1.6.5 accounting for constant sites (-fconst; determined using snp-sites-C) with the built-in model testing (-m MFP) to determine the best phylogenetic model (GTR+F+R2) and 1000 ultrafast bootstraps (-bb 1000) [32]. Pairwise SNP distances were calculated for all pairs of isolates from the SNP alignment using pairsnp v1.0 (https://github.com/gtonkinhill/pairsnp).

To provide a global context for the isolates sequenced in this study, a published clonal complex Eu1 dataset (n = 2,842; Supplementary File 2) spanning fourteen countries was assembled [9, 10, 30, 33–47]. Sequence data was downloaded from the European Nucleotide Archive (ENA) and trimmed using Trimmomatic v0.33 [48]. Sample QC, spoligotype assignment, mapping and phylogenetic tree construction were performed as above. The tree was rooted with a *Mycobacterium caprae* isolate (SRR7617662).

### Transmission Clusters

The R library iGRAPH [49] was used to define putative transmission clusters using a pairwise SNP distance between any two samples of 15 as the threshold. This threshold was chosen as it would allow for the possible identification of older transmission events but also allow for any variance in the rates of mutation amongst the sampled isolates, and has been previously used in a similar analysis of a human *Mycobacterium tuberculosis* dataset [50]. Large clusters were manually divided further on the basis of clear divisions within these clusters observed in the phylogenetic tree (Clusters 5/6 and Clusters 8-12). Transmission clusters with fewer than 50 isolates were not analysed further leaving twelve transmission clusters for further analyses. New alignments were generated for each cluster as described above.

The presence of a temporal signal in each transmission cluster was investigated by plotting the root to tip distance for each isolate, calculated using the R library phytools [51], against its sampling date (Supplementary Figure 1). The slope, x-intercept (most recent common ancestor; MRCA), correlation coefficient and R^2^ value were calculated for each dataset in R. BEAST v1.8.4 [52] was run on each SNP alignment, using tip sampling dates for calibration. Three runs of 10^8^ Markov chain Monte Carlo (MCMC) iterations were performed using a HKY substitution model, strict or constant molecular clock and constant or exponential population size and growth (12 separate runs) for each transmission cluster. The performance of each model was assessed through the comparison of posterior marginal likelihood estimates [53, 54] and the model with the highest Bayes factor [55] (strict clock/constant population size) was selected for each transmission cluster (Supplementary Table 1). The three selected MCMC runs were combined using LogCombiner v1.8.4 (10% burnin) and convergence was assessed (posterior effective sample size (ESS) > 200 for each parameter). A maximum clade creditability tree summarizing the posterior sample of trees in the combined MCMC runs was produced using TreeAnnotator v1.8.4. To confirm the temporal signal in each tree generated, the R library TIPDATINGBEAST [56] was used to resample tip dates from each alignment to generate 20 new datasets with randomly assigned dates. BEAST was then run on each new dataset using the same strict clock priors (Supplementary Figure 2). If the estimated substitution rates in the observed data did not overlap with the estimated substitution rates in the randomized data then the temporal signal observed in the observed data was considered not to be obtained by chance.

Transmission reconstruction was performed on each cluster using the R library TransPhylo [57] which allows for unsampled cases and within-host diversity. The same parameters (gamma shape = 1.6; scale = 3.5) were used for the infection and generation time prior distribution. The TransPhylo algorithm was run three times for 10^7^ MCMC iterations sampling every 200,000 states and a burnin of 10% on each cluster using the MCC trees generated previously. The R library coda [58] was used to assess convergence (Gelman and Rubin’s Convergence Diagnostic < 1.05) and ESS values > 100 for within-host diversity, reproductive rate and sampling proportion (Supplementary Table 2). Post processing of each TransPhylo run was performed in R.

The BEAST2 [59] package BASTA (Bayesian Structured coalescent Approximation) [60] was used to estimate transmission rates between badgers and cattle, defined as demes, in each transmission cluster. A strict clock/equal population size model was used and the BASTA analysis was repeated three times and run for 3 x 10^8^ MCMC iterations with 10% burnin. Convergence was assessed as above. Post processing of the BASTA analysis was performed in R.

SNP-scaled phylogenetic trees were calculated for each transmission cluster using pyjar (https://github.com/simonrharris/pyjar) [61] and plotted using the R libraries treeio and ggtree [62, 63].

### Cattle Movements

bTB metadata was extracted from APHA’s Sam database which records all statutory bTB testing information. Cattle movement metadata was extracted from APHA’s copy of the Department of Environment, Food and Rural Affairs’ (DEFRA)’s Cattle Tracing System (CTS). Movement data were extracted for 727/752 cattle where the ear tag could be matched to the Sam database (it only became a legal requirement to record cattle movement in the CTS after January 2001 so movement data may be missing for the early part of the RBCT). Movements of TB test reactor cattle that were not subjected to laboratory culture and/or sequencing of *M. bovis*, but may have contributed to the spread of infection, were extracted from the CTS using the following criteria: the animals passed through the same location as an animal with a sequenced isolate, the animals were born before 2009 and the animals were classified as “reactors”. Animals were classified as reactors if they had a positive tuberculin test result, had an inconclusive test result but were slaughtered and culture positive for *M. bovis*, were culture positive for *M. bovis* following detection by routine meat inspection at a slaughterhouse, or were culture-negative reactors that led to a breakdown with other tuberculin test positive animals. UK grid coordinates were extracted from the Sam database by matching to location IDs. Past breakdown history was extracted by matching herds using county-parish-holding (CPH) numbers. Where multiple herds had the same CPH number, the active dates of the herds were checked and the individual animal test records were used to identify the correct entries. Short stay locations and locations with missing coordinates were excluded by creating animal records that removed missing locations or stays of fewer than eight days. Where subsequent movements occurred, these were connected to the previous movements to create a continuous record. Where the cattle ear tag IDs of sequenced isolates could not be matched to the database, the CPH was used to identify the final location and coordinates for plotting. The final herd of the animals with sequenced isolates was determined as the location closest to death where the length of stay was at least seven days. The data was queried and extracted from the CTS using PostgreSQL. Pairwise geographic distances between each isolate in kilometres were calculated using the distHaversine function from the R library geosphere [64]. Herd and badger locations were randomly shifted by up to 1 km in the horizontal and vertical planes for plotting using the R libraries maps and mapdata [65].

## Results

### Population structure

A total of 1,442 *M. bovis* isolates from badgers (n = 690) and cattle (n = 752) were sequenced and passed QC; the sites of collection for all 1,442 isolates are shown in Figure 1A. The average number of sequenced isolates per trial area was 144 (range: 81-233) and the ratio of cattle to badger isolates varied from 0.22 (trial area D3) to 4.38 (trial area B2; Table 1). All sequenced isolates were collected between 1999 and 2010. The majority (1437/1442; 99.7%) of the isolates were clonal complex Eu1 whilst the remaining five isolates (all SB0134) belonged to an as yet undefined clonal complex (labelled Unknown7 in Loiseau et al. [27]; Figure 1B).

**Figure 1.**
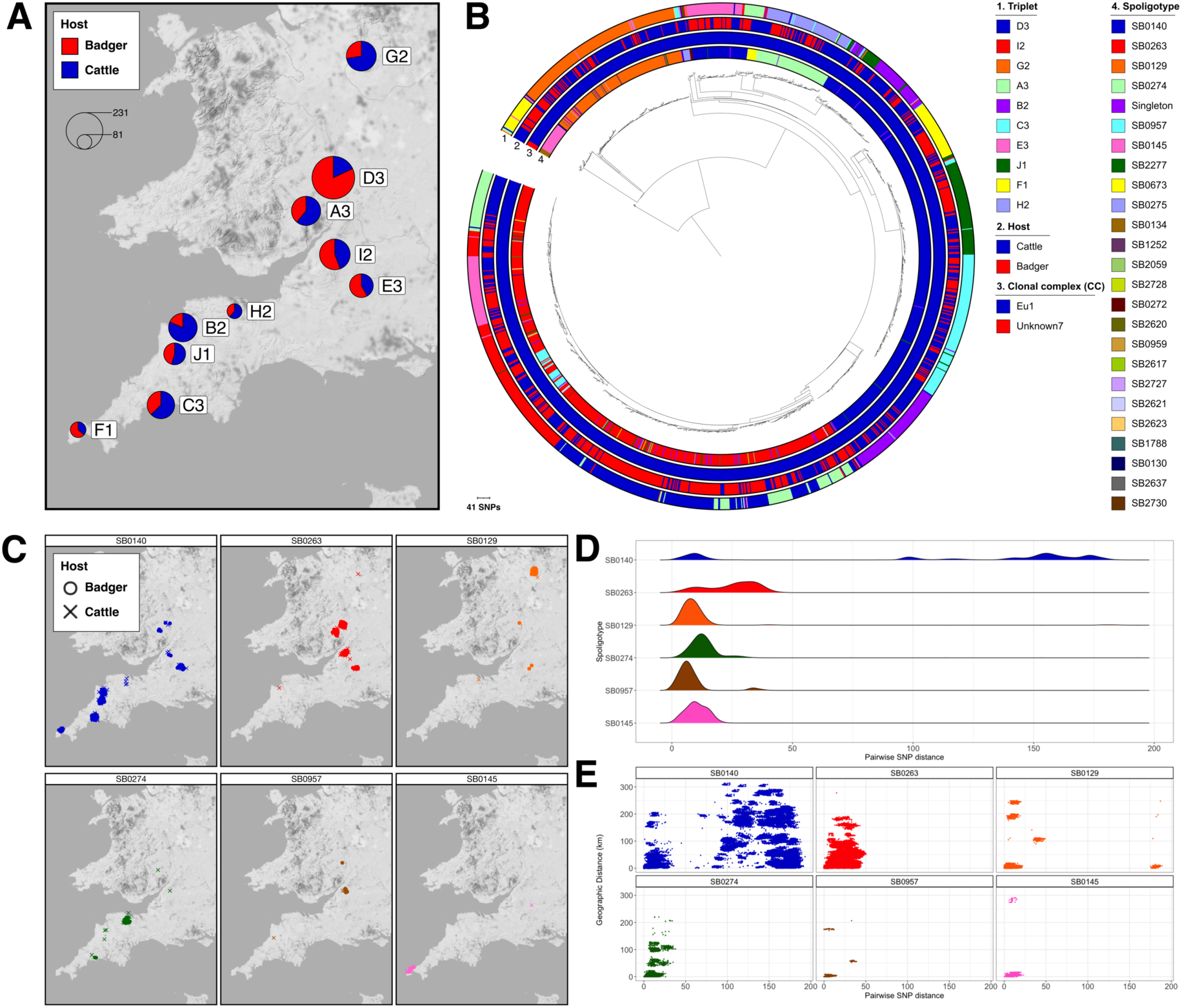
Genomic epidemiology of Randomised Badger Culling Trial (RBCT) dataset: A) Map showing location of isolation for 1,442 sequenced *Mycobacterium bovis* isolates. Isolates collected from badgers and cattle are shown in red and blue respectively. The proportion of samples from each host is shown in the pie charts and the pie charts are scaled according to number of isolates. The RBCT triplet where each of the isolates were collected is labelled; B) Maximum likelihood phylogenetic tree of 1,442 *M. bovis* isolates rooted with isolates from the Unknown7 clonal complex. Trial area, host, clonal complex and spoligotype are shown as datastrips around the outside of the phylogenetic tree; C) Geographical distributions of the six most prevalent spoligotypes in the dataset. The host of each isolate is represented by a different shape: circle for badger and cross for cattle; D) Frequency distributions of pairwise SNP distances between all isolates belonging to the six most prevalent spoligotypes; E) Scatterplots of pairwise SNP distance against geographic distance in kilometres for all pairs of isolates belonging to the six most prevalent spoligotypes.

**Table 1:** Breakdown of the 1,442 sequenced *Mycobacterium bovis* isolates by trial area and host. The table is ordered by total number of isolates from high to low.

| **Trial area** | **Date range** | **Cattle (n)** | **Badgers (n)** | **Total** |
| --- | --- | --- | --- | --- |
| **D3** | 2002-2010 | 42 | 191 | 233 |
| **I2** | 2002-2007 | 74 | 93 | 167 |
| **G2** | 2000-2006 | 118 | 45 | 163 |
| **A3** | 2000-2007 | 97 | 62 | 159 |
| **B2** | 1999-2007 | 127 | 29 | 156 |
| **C3** | 1999-2006 | 94 | 56 | 150 |
| **E3** | 2000-2007 | 54 | 75 | 129 |
| **J1** | 2002-2008 | 65 | 54 | 119 |
| **F1** | 2000-2005 | 31 | 54 | 85 |
| **H2** | 2000-2006 | 50 | 31 | 81 |
| **Total** | **1999-2010** | **752** | **690** | **1442** |

Over 60 unique spoligotypes were identified with the most prevalent being SB0140 (n = 531), SB0263 (n = 491), SB0129 (n = 147), SB0274 (n = 85), SB0957 (n = 34) and SB0145 (n = 32). With the exception of SB0140 and SB0263, which were found in multiple trial areas, the geographical distributions of the most prevalent spoligotypes were largely confined to a single trial area (Figure 1C). Examination of the pairwise SNP distances of isolates within the above spoligotypes showed that there were considerable differences in diversity amongst the spoligotypes (Figure 1D). High levels of diversity were observed in spoligotypes SB0140 and SB0129 reflecting the phylogenetic structure of the isolates with these spoligotypes. Figure 1E shows pairwise SNP distances for all isolates plotted against geographic distance for each of the most prevalent spoligotypes.

### Transmission

#### Transmission clusters

A total of twelve putative transmission clusters, containing 1224/1442 (84.9%) of the isolates, were defined using a conservative threshold of 15 SNPs. The clusters varied in size between 54 (Cluster 2) and 193 (Cluster 9) isolates (Table 2). The ratio of cattle to badger isolates in each transmission cluster varied from 0.15 (Cluster 9) to 5.44 (Cluster 12; Table 2). The phylogenetic tree of all 1,442 isolates with the transmission clusters overlaid on it is shown in Figure 2A and the geographical distribution of each transmission cluster is shown in Figure 2B. The geographical distribution of the transmission clusters was strongly associated with trial area, with the majority of isolates from a transmission cluster found in the same trial area (Figure 2B).

**Figure 2:**
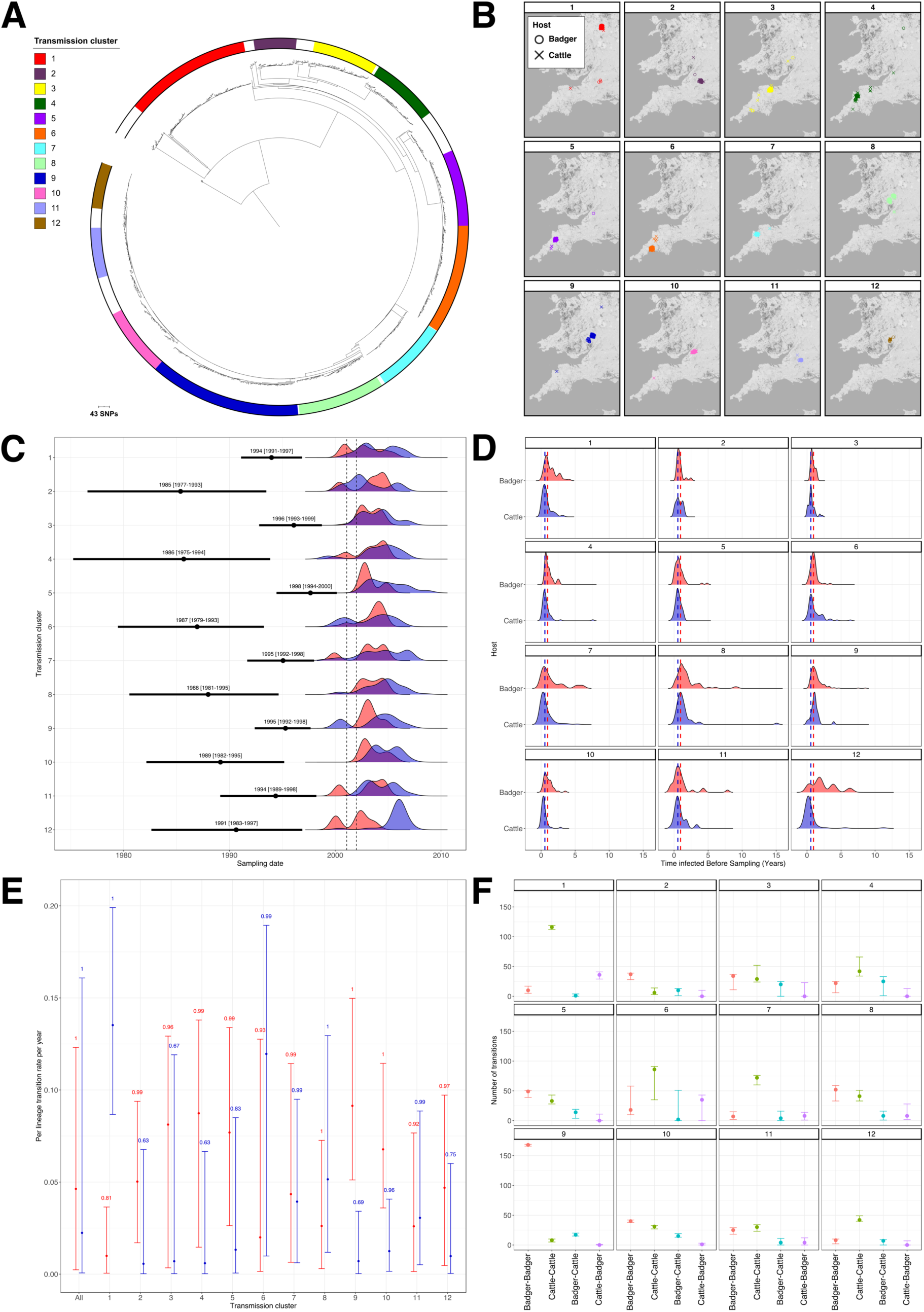
Transmission in the Randomised Badger Culling Trial (RBCT) dataset: A) Maximum likelihood phylogenetic tree of 1,442 isolates with the twelve putative transmission clusters annotated; B) Geographical distributions of the twelve putative transmission clusters. The host of each isolate is represented by a different shape: circle for badger and cross for cattle; C) Molecular dating of transmission clusters. The inferred median and 95% confidence intervals of the MRCA for each transmission cluster is shown in black. The dates of collection of samples within each transmission cluster are shown as frequency distributions and coloured according to host (red for badgers and blue for cattle). The time period of the suspension of badger culling due to FMD is represented by dashed lines; D) Median length of time of infection for all isolates before sampling per transmission cluster. The medians of all isolates are shown by red and blue dashed lines for cattle and badgers respectively; E) Estimated inter-species transmission rates for each transmission cluster. The vertical lines show the lower and upper (2.5% and 97.5%) bounds of the transmission rate distribution for each transmission cluster. The values above the vertical lines represent the posterior probability of each rate and the distributions are coloured according to direction of transmission (red for badger-to-cattle and blue for cattle-to-badger transmission); F) Number of transmissions between known and estimated species counted on each phylogenetic tree in the posterior distribution for each transmission cluster. The vertical lines show the lower and upper (2.5% and 97.5%) bounds of the distributions. The distributions are coloured according to the type of transmission (red for badger-badger, green for cattle-cattle, blue for badger-cattle and purple for cattle-badger).

**Table 2:** Breakdown of the 12 putative transmission clusters by host. The table is ordered by total number of isolates from high to low.

| Cluster | Badgers (n) | Cattle (n) | Total |
| --- | --- | --- | --- |
| Cluster 9 | 168 | 25 | 193 |
| Cluster 1 | 46 | 115 | 161 |
| Cluster 6 | 53 | 86 | 139 |
| Cluster 8 | 61 | 49 | 110 |
| Cluster 5 | 50 | 47 | 97 |
| Cluster 7 | 16 | 76 | 92 |
| Cluster 10 | 41 | 46 | 87 |
| Cluster 4 | 19 | 67 | 86 |
| Cluster 3 | 34 | 49 | 83 |
| Cluster 11 | 30 | 34 | 64 |
| Cluster 12 | 9 | 49 | 58 |
| Cluster 2 | 38 | 16 | 54 |
| Total | 565 | 659 | 1224 |

A multimodal distribution was observed for pairwise SNP distances of isolates assigned to each of the transmission clusters (Supplementary Figure 3). The first mode comprised pairwise differences of 400 - 500 SNPs and was made up of comparisons of isolates from Eu1 clades deeper in the phylogeny, and the second and third modes between 100-200 SNPs were comprised of isolates from more closely related clades. The final modes between 0 and 50 SNPs were made up of comparisons of isolates from the same clade and here the within and between transmission cluster comparisons overlapped, although there was a clear peak below 15 SNPs representing the transmission clusters themselves. There were no observable differences between the distributions when the host of each isolate in a pairwise comparison was considered i.e. there were no host-specific patterns of genetic relatedness.

#### Temporal analyses of transmission clusters

To describe the temporal dynamics of the transmission clusters, each cluster was independently tested for evidence of temporal signal. Comparison of root to tip distances with sampling dates did not find significant correlations for any of the transmission clusters (Supplementary Figure 2). However, dated tip randomisation (DTR) analyses, where evidence of a temporal signal is shown by a lack of overlap between the estimated substitution rates of the observed data and the randomised datasets, showed that there was no overlap between the highest posterior densities (HPD) of the real and randomised datasets for 5/12 of the transmission clusters (Supplementary Figure 3). In a further 5/12 there were overlaps between the HPDs but not medians of the real dataset and one or more of the randomised datasets. For the final two clusters (Cluster 2 and Cluster 4), the median substitution rates of one and five randomised datasets respectively overlapped that of the real datasets (Supplementary Figure 3).

The median substitution rate of each transmission cluster varied between 0.51 (Cluster 12) and 6.0 (Cluster 3) substitutions per genome per year (Supplementary Table 3). Phylogenetic dating analysis using BEAST showed that the estimated date of the MRCA of each transmission cluster varied between 1985 (Cluster 2; 95% confidence interval [CI]: 1977 to 1993) and 1997 (Cluster 5; 95% CI: 1994 to 2000; Figure 2C). The median difference between the MRCA and the date of collection of the first sample was 8.1 (range 4.0 – 15.0) years.

#### Transmission analysis with TransPhylo

TransPhylo was used to estimate the number of unsampled cases. The median number of sampled cases per transmission cluster was 89.5 (range: 54 - 193) compared to a median of 130.1 (range: 41 - 203) inferred unsampled cases suggesting a median case finding rate of 43.2% (range: 28.3% - 74.2%; Supplementary Figure 4). The median inferred time from infection to sampling across all transmission clusters was 0.56 years (95% CI: 0.07 – 2.26 years) for cattle and 0.96 years (95% CI: 0.25 – 3.36 years) for badgers (Figure 2D).

A total of 84 highly supported transmission pairs (posterior probability of transmission between isolate 1 and isolate 2 > 0.5) were identified within the twelve transmission clusters (Table 3) using TransPhylo. The majority of these transmissions (60/84) were within-species whilst 24/84 were between species. No highly supported transmission pairs were identified in Cluster 3. The median pairwise SNP distances for the highly supported transmission pairs across all transmission clusters were 1 (range: 0-8), 1 (range: 0-5) and 1 (range: 0-6) for cattle to cattle, badger to badger and between-species transmission respectively.

**Table 3:** Highly supported transmission pairs within each transmission cluster. For each transmission cluster, the number of intra-and inter-species transmission pairs with a probability > 0.5 is listed. The median SNP distance for each set of transmission pairs is given in parentheses.

| **Transmission cluster** | **Number highly supported transmission pairs** | **Number Cattle-Cattle transmission pairs (median SNP distance)** | **Number Badger-Badger transmission pairs (median SNP distance)** | **Number Between-species transmission pairs (median SNP distance)** |
| --- | --- | --- | --- | --- |
| **Cluster 1** | 9 | 4 (2) | 1 (0) | 4 (1) |
| **Cluster 2** | 1 | 0 | 0 | 1 (1) |
| **Cluster 3** | 0 | 0 | 0 | 0 |
| **Cluster 4** | 1 | 1 (1) | 0 | 0 |
| **Cluster 5** | 14 | 3 (1) | 7 (1) | 4 (1) |
| **Cluster 6** | 5 | 4 (0.5) | 1 (2) | 0 |
| **Cluster 7** | 15 | 11 (1) | 0 | 4 (1) |
| **Cluster 8** | 10 | 4 (1) | 3 (1) | 3 (1) |
| **Cluster 9** | 17 | 0 | 12 (0) | 5 (3) |
| **Cluster 10** | 2 | 1 (1) | 1 (0) | 0 |
| **Cluster 11** | 5 | 3 (2) | 1 (2) | 1 (5) |
| **Cluster 12** | 5 | 0 | 3 (7) | 2 (4) |
| **Total** | **84** | **31** | **29** | **24** |

#### Directionality of transmission between host species

BASTA was used to determine the dominant direction of transmission between host species and found higher rates of transmission from badgers to cattle in 8/12 transmission clusters and from cattle to badgers in 4/12 clusters (the confidence intervals for each direction overlapped in all clusters except clusters 1 and 9; Figure 2E). For the networks with higher badger to cattle transmission, this direction of infection occurred between 1.1 (Cluster 7) and 14.8 (Cluster 4) times more frequently than in the opposite direction. By comparison, in the four networks with higher cattle to badger transmission, the frequency was between 1.2 (Cluster 11) and 13.6 (Cluster 1) times higher than in the opposite direction. The overall median badger to cattle transmission rate for all transmission clusters was 2.1 (95% CI: 0.8-3.8) times higher than the cattle to badger transmission rate.

As BASTA does not directly calculate the transmission rate within-species, the lower bound of the number of transmissions (the count of transitions in the posterior trees) between different animals, regardless of host, was also calculated for each transmission cluster (Figure 2F). For each of the clusters, the estimated number of transmission events is consistent with the estimated inter-species transmission rates. Across the twelve clusters, the number of within-species transmission events was higher than the between-species transmissions with the average number of cattle to cattle transmission events 4.9 (range 0-31) times greater than the number of cattle to badger transmission events and 17 (range 0.5-116) times greater than the number of badger to cattle transmission events. The number of badger to badger transmission events was 4.7 (range 0.9-10) times higher than the number of badger to cattle transmission events and 4.5 (range 0-40) times higher than the number of cattle to badger transmission events.

#### Other Transmission dynamics

Metadata from the SAM database was incorporated for each of the cattle within a transmission cluster that could be matched, which allowed examples of recurrence (detection of infections subsequent to previous outbreaks or breakdowns), superspreading (individual hosts that have a disproportionate effect on the spread of infection) and long-distance transmission (transmission between different trial areas) to be characterised.

#### Recurrence

A total of 47 isolates formed a distinct clade within the Cluster 6 phylogeny (Figure 3B). Of these, 25 were isolates from cattle slaughtered between February 2001 and October 2005 as part of three recorded breakdowns on the same farm comprising 35 confirmed cases (Figure 3A). Based on the date of slaughter, these isolates were divided into six slaughter groups (Figure 3A). Two pairs of isolates from different animals (1 and 20; 6 and 23) and different slaughter groups (1 and 5) were 0 SNPs apart despite the subsequently infected animals (20 and 23) only moving to the farm a year after the first animals were slaughtered (Figure 3A). The subsequently infected animals were then slaughtered three years later as part of a later breakdown. The majority of the rest of the cattle isolates in this clade were also very similar despite the long time periods from when these animals arrived on the farm and when they were slaughtered. Cattle were still present on the farm between the different breakdowns suggesting that infection was being maintained locally, either in this or a neighbouring herd or else within the local badger population.

**Figure 3:**
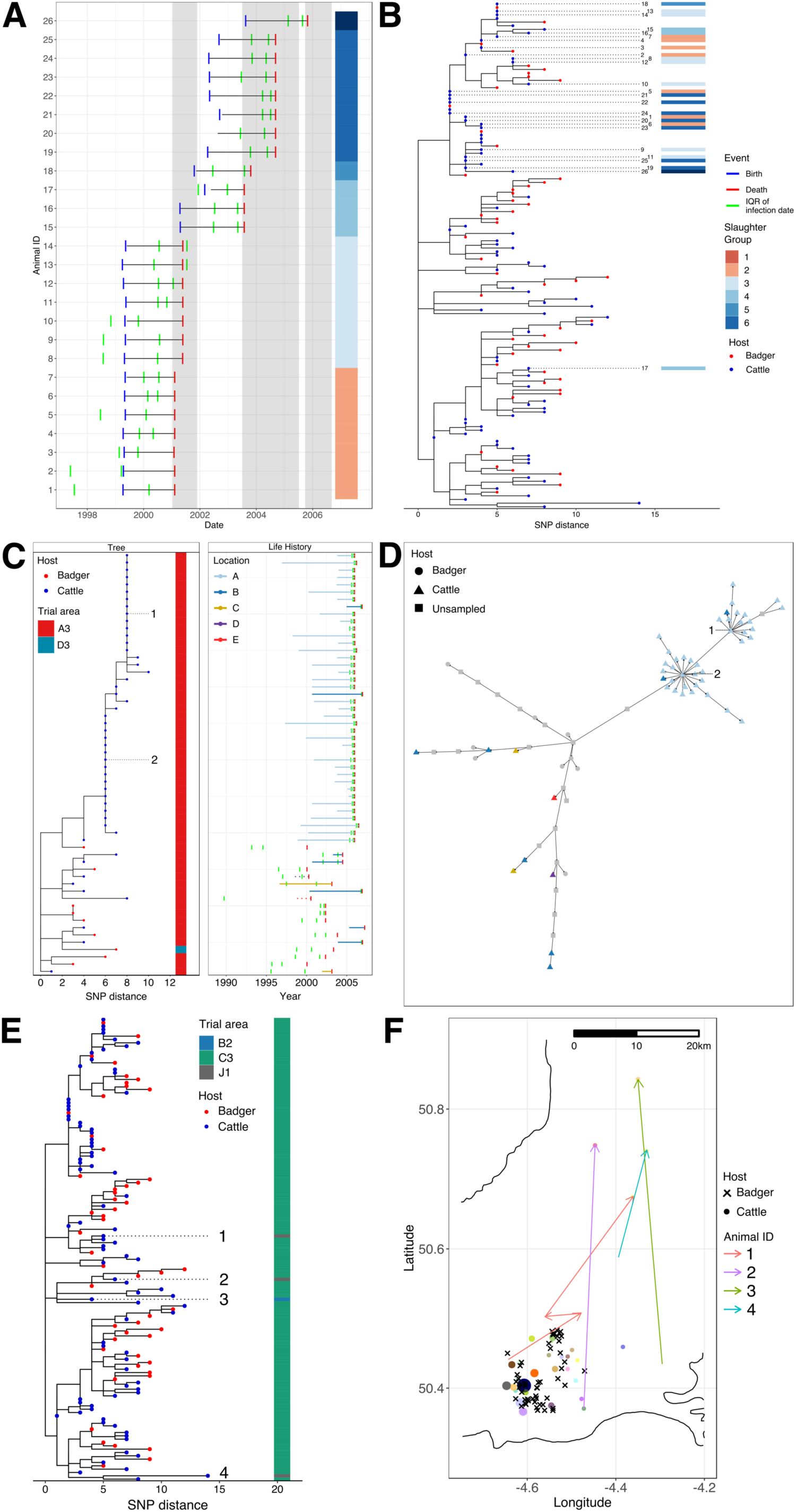
Integration of genomics and cattle movement data: A) Life histories of animals included in Cluster 6. Date of birth is shown in blue, date of death in red and the interquartile range (IQR) for the estimated date of infection is shown in green. The grey shading represents breakdowns and Slaughter Groups are labelled based on date of death; B) SNP-scaled phylogenetic subtree for Cluster 6. Tips are coloured according to host (red for badger and blue for cattle), relevant isolates are labelled and Slaughter Group is shown as a data strip; C) SNP-scaled phylogenetic tree for Cluster 12. Tips are coloured according to host (red for badger and blue for cattle), isolates inferred as superspreaders are labelled and Trial area is shown as a data strip. Life histories are shown for each isolate. Date of birth is shown in blue, date of death in red and the interquartile range for the estimated date of infection is shown in green. The farm location identifier of each cattle isolate is coloured according to the legend and the length of each bar reflects the length of time an animal spent at its final location; D) Transmission network for Cluster 12. The shape of each node is based on the host (circle for badger or triangle for cattle) or else a square for inferred unsampled cases. Nodes are coloured based on their location (grey for unsampled and badger isolates which don’t have a farm location identifier). Isolates inferred as superspreaders are labelled; E) SNP-scaled phylogenetic tree for Cluster 6. Tips are coloured according to host (red for badger and blue for cattle), isolates highlighted as examples of long-distance transmission are labelled and the Trial area for each isolate is shown as a data strip; F) Map showing the location of isolation for each isolate in Cluster 6. The shape of each isolate is based on the host (cross for badger or circle for cattle), the colour of the cattle isolates is based on the farm location and the size of the cattle isolates reflects the number of animals in Cluster 6 at that location. The movements of the animals from which isolates highlighted as examples of long-distance transmission were collected are shown as arrows and coloured according to animal identifier. Herd and badger locations were randomly shifted by up to 1 km in the horizontal and vertical planes.

#### Superspreading

The structure of the Cluster 12 phylogeny showed a number of cattle isolates clustering together within a very flat tree structure i.e. the majority of these isolates were 0 SNPs apart (Figure 3C). Of these, 38 isolates were from animals slaughtered at a single location (A) as part of a breakdown between April 2005 and July 2007 that identified 59 reactors from which 39 had *M. bovis* cultured. The resulting transmission network inferred two distinct superspreading events of 13 and 21 cases inferred to be centred around two animals (1 and 2; Figure 3D). Two of the animals in these superspreading events were from a different location (B) though this was only 0.83 km away, suggesting potential epidemiological links between locations A and B.

#### Long-distance transmission

Isolates from four cattle in Cluster 6 were not from the predominant trial area (C3) of this cluster (Figure 3E). For three of these isolates (1, 2 and 3), the inferred dates of infection showed that the animals, which had moved between farms, were likely infected at a location in or near trial area C3. For the remaining animal, 4, the inferred date of infection didn’t overlap with a previous location in or near trial area C3 but the location data for 58 days of this animal’s life is missing from the database. The distances moved by the infected animals ranged between 23 and 46 km providing evidence of movement mediated transmission (Figure 3F).

## Discussion

The RBCT was set up to assess the impact of badger culling on the incidence of bTB in nearby cattle herds. In this study, the resulting badger and cattle isolates have been sequenced, meaning that, for the first time, WGS of large numbers of *M. bovis* isolates from co-located populations of both species collected contemporaneously in well-defined geographical areas can be used to address key questions surrounding bTB transmission in the high-risk regions of England.

A total of 60 unique spoligotype patterns were identified in the dataset though the majority of the isolates (1320/1442; 91.5%) were made up of the six most prevalent spoligotypes confirming that there is relatively little genetic heterogeneity at this level amongst *M. bovis* in the high prevalence areas of England. The observed prevalence of spoligotypes in this study closely matched previously published data, generated using traditional typing methods, from the same time period [66], showing that the dataset accurately reflects the known population structure of *M. bovis* at that time.

Genotyping methods such as spoligotyping are used to infer close relationships between isolates (low genetic diversity) and assume monophyly. However, it is clear from plotting spoligotypes on the phylogenetic tree in Figure 1B, that some of the spoligotypes, in particular SB0140, are polyphyletic. Whilst the majority of SB0140 isolates sit adjacent to each other in the phylogenetic tree, there is a single clade that sits separately with the SB0274 and SB0673 clades falling between them. Another example is the intermingling of SB0957 and SB0263 isolates in trial area I2. An alternative way to view this data is to calculate and plot pairwise SNP distances for each of the most prevalent spoligotypes (Figure 1D). From this, we see that while there is a maximum of approximately 50 SNPs between members of four of the six most prevalent spoligotypes, for SB0140 and SB0129 the maximum pairwise SNP distance was between 150 and 200 SNPs, demonstrating higher levels of diversity for the *M. bovis* population as evidenced from genome-wide data compared to traditional typing data.

As we observed in our data, it has been recognised for a long time that spoligotypes can be homoplasic and identical spoligotype patterns can be found in phylogenetically unrelated strains [66]. Despite this, spoligotyping, along with Mycobacterial interspersed repetitive unit-variable number tandem repeat (MIRU-VNTR) analysis, has continued to be the most commonly applied method for genotyping *M. bovis* as it is cheap and comparatively straightforward to implement in the laboratory. However, given the much higher resolution offered by WGS and the fact that national bodies such as APHA and the United States Department of Agriculture (USDA) are now moving towards routinely sequencing all cases of *M.bovis*, it may now be time to move towards a SNP-based method of typing *M. bovis* isolates similar to the Coll method adopted for typing *M. tuberculosis sensu stricto* [67].

The predominance of clonal complex Eu1 in our dataset was unsurprising as previous work, including the study that first defined Eu1 [24], has shown that this clonal complex is ubiquitous in Great Britain, Northern Ireland and the Republic of Ireland whilst being uncommon in mainland Europe, where *M. bovis* genetic diversity is much greater [42]. Previous work using both PCR and genomics to assign clonal complex shows that Eu1 is likely the most predominant globally-circulating *M. bovis* lineage and it has been found in many countries that have historically traded cattle with the UK such as New Zealand, the USA, Mexico and Uruguay [33–35, 41]. The presence of this clonal complex and its spoligotypes such as SB0140 and SB0263 in these countries suggests that Eu1 has been present in the UK for at least 200 years or more. This is supported by the high level of pairwise SNP diversity observed within SB0140.

The Unknown7 isolates in the dataset were found in trial areas A3, D3 and G2 and were a maximum of 12 SNPs apart. The small number of isolates suggest that it is uncommon in the UK and the low level of genetic diversity suggests that it may be part of a single, recent introduction. This clonal complex has also been found in France, Mali and the USA [27, 33, 42] and given the geographical proximity of France and the ongoing trade of cattle between the two countries this may be the most likely origin for this lineage.

The prevailing hypothesis for the population history of *M. bovis*, in particular Eu1, in the UK is that there was a single introduction followed by long term endemicity and a population bottleneck due to effective control measures beginning in the 1930s [66]. However, the observed phylogenetic structure of isolates when incorporated with a global collection of Eu1 isolates, indicates that there have likely been multiple, perhaps as many as four, introductions of Eu1 into England (Supplementary Figure 5). Whilst we did not attempt to date these introductions in this study, the availability of archived isolates from the 1980s along with contemporaneous isolates will be used alongside the RBCT dataset and other published UK datasets in future work to provide estimated dates for these introductions.

Due to the large size and clear phylogenetic and geographical structure of the dataset as well as our study aims, we defined transmission clusters using a conservative pairwise SNP threshold of 15 SNPs. There is currently no consensus as to what is the best threshold to apply to *Mycobacterium* genome datasets with previous studies using thresholds between three and fifteen SNPs [40, 50]. We chose the 15 SNP threshold as it would allow for the possible identification of older transmission events but also allow for any variance in the rates of mutation amongst the sampled isolates. We chose to use the software package TransPhylo as it integrates dates of isolate collection and genetic relatedness and allows for within-host diversity and unsampled cases. The first step of the analysis was to generate molecular dated phylogenies for each of the transmission clusters. Assessing the presence of temporal signal in a genome dataset is typically done in two ways: examining the linear relationship between root to tip distance and sampling date (under a perfect clocklike behaviour, then R^2^ = 1 [68]), and dated tip randomization (DTR) analysis. In DTR, the dates of sampling are repeatedly shuffled amongst the taxa and the clock rates between the observed and random data calculated and compared [69]. If there is no overlap between the estimated substitution rates of the observed data and the randomized datasets then we can conclude that the observed dataset has a stronger temporal signal than expected by chance [69]. We obtained very low or negative values for R^2^ for all our transmission clusters which is normally interpreted as evidence for a lack of temporal signal or else overdispersion in lineage-specific clock rates [70] (Supplementary Figure 2). The previously reported slow substitution rate of *M. bovis* [10, 34] and the short window of sampling (twelve years) may explain the lack of association between root to tip distances and sampling dates. As root to tip regression is only a tool for exploratory analysis [70] we performed DTR on all of the transmission clusters (Supplementary Figure 3). From this, we observed that there was strong evidence of temporal signal in 5/12 of the transmission clusters, moderate temporal signal in 5/12 transmission clusters and weak temporal signal in two clusters, particularly Cluster 2 (Supplementary Figure 2). Despite the especially weak evidence of temporal signal in Cluster 2, we decided to include it in our analyses as, given the close relatedness of all of the clusters, it is highly likely that the mutational process, and therefore the molecular clock, will be similar in each of them. As well as being used as input for TransPhylo molecular dating of each of the transmission clusters provided additional insights into the dataset.

Firstly, we were able to calculate substitution rates for each transmission cluster which ranged between 0.5 and 6 SNPs per genome per year (Supplementary Table 3). Published estimates for the median substitution rate of *M. bovis* vary between 0.15 and 0.53 substitutions per genome per year [10, 34]. Previous work has shown that there are lineage and study specific differences in the substitution rates within the *Mycobacterium tuberculosis* complex (MTBC) [71]. This analysis showed that higher substitution rates were found in smaller datasets with narrow sample date ranges, which may explain the much higher substitution rates of up to six substitutions per genome per year we observed in our analyses. Secondly, we were able to infer the date of the MRCA of each transmission cluster (Figure 2C). The short time period (4 – 15 years) between the inferred date of the MRCAs and the earliest sample collection dates suggests that the transmission clusters are likely the result of recent seeding events and not a consequence of endemic disease in the form of long-term maintenance within herds or an endemic wildlife reservoir, in this case badgers. The introduction of a compulsory test and slaughter scheme in the UK in the 1950s saw a sustained decline in the annual number of infected animals removed as TB test reactors and infected cattle herds with only a few hundred reactors being detected annually in the early 1980s [66]. However, this decline was reversed from the late 1980s, with the UK now having one of the highest incidences of bTB in Europe. From our analyses, it is clear that the dates for the MRCA of each of the transmission clusters overlap with this population expansion.

From the TransPhylo analysis we were able to estimate what proportion of infected hosts we managed to sample for each transmission cluster (Supplementary Figure 4). Sampling of all hosts infected with a disease is never complete due to a range of factors such as detection, failure to culture and in the case of genomics, issues related to sequence quality. In this study we estimate that we managed to sequence a median of 43.2% of cases across the transmission clusters though this varied from less than 30% to as high as 75% depending on the cluster. Obviously, the success of sampling has an impact on the types and quality of the inferences we are able to make. For instance, we were able to confidently identify and confirm a superspreading event in Cluster 12 due to having sequenced 38/39 of the confirmed cases in a breakdown (see below).

The incubation period of TB in cattle is generally believed to be several months and potentially years, although there is some evidence of much shorter incubation periods in other mammalian species such as cats [72]. There is also typically a lag period (occult period) between infection and detection where infections are undetectable to the standard tuberculin test [73]. The organism may also persist for several years within infected animals before they are detected (latency) and reactivation has not been demonstrated in cattle. To date, there are no firm estimates for either the duration of the occult period or of epidemiological latency, which is problematic for fitting transmission models [74] and predicting the impact of control polices [75]. Based on our analysis using TransPhylo, we can provide estimates for how long both badgers and cattle were infected before sampling. The analysis showed that, on average, badgers were infected for twice as long as cattle before sampling. The median period of infection for cattle of 0.56 years is consistent with the annual testing schedule imposed on cattle during the RBCT. Whilst there was a wide range of estimates for the length of infection the 95% confidence intervals for both badgers and cattle were within the normal lifespans of both species.

We were able to identify a small number of highly supported direct transmission events, defined as transmission pairs that had a posterior probability greater than 0.5 (Table 3). Although the majority (60/84) of these transmission events occurred between the same species, there were also 24 interspecies transmission pairs across the transmission clusters with pairwise SNP distances varying between 1 and 5 SNPs. To date, there is limited evidence of badgers and cattle directly interacting and the majority of transmission is considered to be indirect i.e. through the environment [76]. Given the inferred number of unsampled cases and small number of highly supported transmission pairs, more intensive sampling would need to be performed to better establish transmission dynamics between the different bTB host species. Despite the logistical challenges around detecting and culturing *M. bovis* in environmental samples, the inclusion of samples from faeces and feed troughs and other potential hosts such as rodents and cervids should be an integral part of any future work.

One of the aims of this study was to assess and quantify the directionality of transmission between cattle and badgers. For this we used a Bayesian evolutionary tool, BASTA (Bayesian Structured coalescent Approximation), to estimate the interspecies transmission rates in each of the transmission clusters. BASTA was designed to estimate evolutionary dynamics in structured populations and account for sampling biases. For the majority of our transmission clusters, badger to cattle transmission occurred more frequently even in clusters with approximately equal numbers of cattle and badgers (Figure 2E). It is worth noting that the estimated transmission rates were very low with the median number of badger to cattle transmissions across all transmission clusters estimated as 0.05 transmissions per lineage per year and the median number of cattle to badger transmissions estimated as 0.02 transmissions per lineage per year (Figure 2E). Whilst BASTA does not directly estimate intra-species transmission rates we could calculate the number of transmission events between each host species from the posterior log and tree files. These are conservative counts of the minimum number of transitions between sampled animals and their ancestors but do allow us to compare the number of inter-and intra-species transmissions. From this we were able to demonstrate that inter-species transmission occurs much less frequently than intra-species transmission in our transmission clusters and cattle to cattle transmission is more common than badger to badger transmission (Figure 2F). Three previous studies, each on small geographically localized populations, have used BASTA to estimate rates of transmission between badgers and cattle; the first estimated that badger to cattle transmission was 10.4 times more frequent than cattle to badger transmission, the second estimated that cattle to badger transmission was at least an order of magnitude higher than badger to cattle transmission and the third estimated that cattle to badger transmission was at least three times higher than badger to cattle transmission (a similar result was obtained using a similar transmission analysis package, MASCOT) [9, 12, 13]. These results, along with those described in this study, suggest that the directionality of transmission may vary between sampling area although badger to cattle transmission does appear to be more frequent. What is consistent across all the studies, however, is that intra-species transmission occurs much more frequently than inter-species transmission.

Beyond the original aims of the project such as characterising the population structure of *M. bovis* isolates collected as part of the RBCT and investigating interspecies transmission, the utility of WGS was also shown through its application to other important aspects of bTB transmission in the UK. The combination of genomics and the extensive cattle tracing database allowed us to characterise examples of recurrence, superspreading and long-distance transmission within the dataset. Previous work has shown that prior history of bTB within a herd is an important predictor of breakdown: 38% of herds that clear movement restrictions experience another breakdown within 24 months [77]. This suggests that infection is being maintained within herds despite repeated testing and it is estimated that between 24% and 50% of recurrent breakdowns are due to persistence within the herd [74]. We were able to use pairwise genome comparisons to identify near identical isolates that were collected up to four years apart and which were part of confirmed herd breakdowns. Examination of the cattle movements confirmed that some of these isolates were collected from animals that arrived subsequent to the dates of slaughter of infected animals as part of previous breakdowns. The similarity of these more recent isolates to the earlier isolates would suggest that the animals were infected after their arrival in the new location and that control measures following the prior breakdowns were insufficiently effective.

TransPhylo allowed us to generate plausible transmission networks where star like nodes representative of potential superspreaders (individual hosts that have a disproportionate effect on the spread of infection) could be identified (Figure 3C/3D). We were then able to incorporate data from the CTS to identify the cattle likely acting as the source of the infections. Whilst previous work using modelling or network analysis has highlighted the importance of small numbers of farms or herds as hubs of transmission which act as superspreaders of infection [78, 79], we provide the first evidence, based on genomics and cattle movement data, that particular animals within herds may also act as superspreaders potentially contributing to increased transmission between different locations if these animals are not identified before being moved. We were unable to identify any superspreaders amongst any of the sampled badgers.

From the temporal analysis of the transmission clusters we showed that these clusters are comparatively young and likely recently seeded. The most likely mechanism for this is the movement of infected cattle into a location followed by subsequent onward transmission within the herd and into the local badger populations. Given the median estimate of the MRCA of the transmission clusters was eight years before sampling began, this precluded any possibility of us sampling the index case for any of the transmission clusters. However, by incorporating cattle movement information with our transmission clusters, we were able to identify cattle infected with a particular lineage in one trial area moving to a trial area further away, highlighting the potential for long distance transmission events to seed new transmission clusters (Figure 3E/3F). This was also recently demonstrated by Rossi et al. who identified an imported infected animal or animals as being responsible for a bTB outbreak in a region of England with no previously known wildlife infections [12]. This has important implications for infection control; even with the limited sampling we conducted, the combination of genomics and cattle movements still allowed us to identify these potential seeding events. More targeted testing and sequencing before animals are moved, particularly to lower incidence areas, would potentially identify these likely sources of infection before they are able to become established in other locations.

Potential limitations of our analysis were the choice and number of samples included in the study and known issues surrounding the lack of a strong temporal signal in *M. bovis* that may affect the results of any analyses based on molecular dating. Any sampling strategy we selected would not have been perfect; ideally, we would have tried to sequence all samples collected as part of the RBCT; however, this was not possible due to cost and manpower constraints so we chose to sequence only the badger and cattle isolates collected from proactive triplets excluding isolates from infected badgers culled in reactive triplets and infected cattle culled as part of contemporaneous breakdowns. From our TransPhylo analysis we estimated that we managed to sample approximately 40% of infected cases across our transmission clusters. Despite this, the size of the dataset was still large enough to generate several large transmission clusters that allowed us to draw robust conclusions about transmission, notably directionality of transmission between badgers and cattle. Comparison of the spoligotype distribution in our study to earlier work confirmed that our dataset was representative of the known population structure during the RBCT.

We know from previous work that the lack of a strong temporal signal is a potential issue when attempting to accurately date the origin of particular lineages [71]. The results of the dated tip randomization analysis indicated that there was moderate or strong temporal signal in nearly all of our transmission clusters; however, two of our transmission clusters notably Cluster 2 had a weak temporal signal. The range of substitution rates we estimated for some of our transmission clusters was also higher than previously observed which may have affected the estimated dates of those transmission cluster’s MRCAs. Overall, however, even if individual networks such as Cluster 2 with little or no temporal signal or Cluster 3 with a high substitution rate are of concern, the conclusions we have drawn are based on considering the results from twelve different transmission clusters composed of over 1,200 genomes and thus can be considered robust.

Multiple previous studies have shown that bTB transmission is complicated, unlikely to be driven by a single mechanism and is strongly associated with the setting and host dynamics of the system being studied. Here we used the largest single country genome dataset alongside the national cattle movement database to attempt to address key questions around bTB transmission in a multi-host, intensive setting. Whilst both the TransPhylo and BASTA results support inter-species transmission with some evidence that there is broadly more badger to cattle transmission than in the opposite direction, it is clear that the majority of ongoing transmission is occurring within cattle herds and within the badger populations. Spillover in either direction could then be considered to be occurring at a low level and, based on the dates of their MRCAs, the transmission clusters we defined are likely to have been the result of recent seeding events and are primarily being maintained by within-species transmission. We have also provided the first genomics-based estimates for the length of time that badgers and cattle are infected with bTB before sampling. Finally, we were able to characterise recurrence, superspreading and long-distance transmission within our transmission clusters.

## Data availability

Raw sequencing reads were deposited at the European Nucleotide Archive (https://www.ebi.ac.uk/ena/browser/home) under project PRJEB19799; all accessions used in this project are listed in Supplementary File 1. Metadata for the sequenced isolates is available on pubMLST (https://pubmlst.org/projects/mbovis-eradbtb).

## Code availability

The R code used to perform data analyses in this study is available in GitHub (https://github.com/avantonder/RBCT).

## Acknowledgements

This work was supported by a joint Biotechnology and Biological Sciences Research Council (BBSRC) research grant under grant number BB/N00468X/1 and Department for Environment Food and Rural Affairs (Defra) research project SE3298. Glyn Hewinson holds a Sêr Cymru II Research Chair funded by the European Research Development Fund and Welsh Government. Thanks to Matt Mayho and Richard Ellis for providing advice on Illumina library construction, Mathew Beale for fruitful discussions and feedback regarding the ongoing analysis and manuscript and, Adrian Whatmore for coordinating Defra research project SE3298.

## Author contributions

A.J.K.C., R.G.H., J.L.N.W. and J.P. conceived the study. J.D. cultured, heat inactivated and submitted the *M. bovis* isolates for sequencing. L.G. and A.P.M. extracted metadata for the study isolates from the CTS and Sam databases and uploaded this metadata to a BIGSdb database created by K.A.J. A.J.v.T. and M.T. performed the data analysis. A.J.v.T. coordinated the study and wrote the initial draft of the manuscript. A.J.v.T, A.J.K.C., E.P. P.J.H., J.L.N.W. and J.P. contributed to the final version of the manuscript. All authors read and approved the manuscript.

## Supplementary Tables and Figures

**Supplementary Table 1:** Model performance based on Maximum Likelihood Estimates (MLE) and Bayes Factors for all transmission clusters

| **Transmission cluster** | **Model** | **log MLE** | **log Bayes Factor** | **Strength of Evidence (Kass & Raftery, 1995)** |
| --- | --- | --- | --- | --- |
| **Cluster 1** | Relaxed exponential | -5536634 | 0.056 | Not worth more than a bare mention |
|  | Relaxed constant | -5536655 | 21.289 | Very strong |
|  | Strict exponential | -5536634 | - | - |
|  | **Strict constant** | **-5536659** | **25.277** | **Very strong** |
| **Cluster 2** | Relaxed exponential | -5534141 | - | - |
|  | Relaxed constant | -5534156 | 14.906 | Very strong |
|  | Strict exponential | -5534146 | 4.800 | Strong |
|  | **Strict constant** | **-5534161** | **19.609** | **Very strong** |
| **Cluster 3** | Relaxed exponential | -5535466 | - | - |
|  | Relaxed constant | -5535498 | 31.000 | Very strong |
|  | Strict exponential | -5535468 | 2.014 | Positive |
|  | **Strict constant** | **-5535505** | **39.717** | **Very strong** |
| **Cluster 4** | Relaxed exponential | -5534382 | 0.101 | Not worth more than a bare mention |
|  | Relaxed constant | -5534394 | 12.002 | Very strong |
|  | Strict exponential | -5534382 | - | - |
|  | **Strict constant** | **-5534396** | **13.793** | **Very strong** |
| **Cluster 5** | Relaxed exponential | -5534469 | - | - |
|  | Relaxed constant | -5534474 | 4.936 | Strong |
|  | Strict exponential | -5534470 | 1.661 | Positive |
|  | **Strict constant** | **-5534476** | **7.052** | **Very strong** |
| **Cluster 6** | Relaxed exponential | -5535426 | - | - |
|  | Relaxed constant | -5535452 | 25.594 | Very strong |
|  | Strict exponential | -5535428 | 2.099 | Positive |
|  | **Strict constant** | **-5535453** | **27.358** | **Very strong** |
| **Cluster 7** | Relaxed exponential | -5533415 | - | - |
|  | Relaxed constant | -5533417 | 2.257 | Positive |
|  | Strict exponential | -5533421 | 5.893 | Very strong |
|  | **Strict constant** | **-5533421** | **6.083** | **Very strong** |
| **Cluster 8** | Relaxed exponential | -5535839 | - | - |
|  | Relaxed constant | -5535863 | 23.890 | Very strong |
|  | Strict exponential | -5535841 | 2.074 | Positive |
|  | **Strict constant** | **-5535866** | **26.912** | **Very strong** |
| **Cluster 9** | Relaxed exponential | -5538088 | - | - |
|  | Relaxed constant | -5538147 | 59.427 | Very strong |
|  | Strict exponential | -5538088 | 0.389 | Not worth more than a bare mention |
|  | **Strict constant** | **-5538157** | **69.010** | **Very strong** |
| **Cluster 10** | Relaxed exponential | -5534887 | - | - |
|  | Relaxed constant | -5534891 | 3.526 | Strong |
|  | Strict exponential | -5534888 | 0.799 | Not worth more than a bare mention |
|  | **Strict constant** | **-5534894** | **7.076** | **Very strong** |
| **Cluster 11** | Relaxed exponential | -5533358 | 1.383 | Positive |
|  | Relaxed constant | -5533358 | 0.961 | Not worth more than a bare mention |
|  | Strict exponential | -5533357 | - | - |
|  | **Strict constant** | **-5533361** | **3.610** | **Strong** |
| **Cluster 12** | Relaxed exponential | -5533082 | 1.546 | Positive |
|  | Relaxed constant | -5533081 | - | - |
|  | **Strict exponential** | **-5533085** | **4.393** | **Strong** |
|  | Strict constant | -5533083 | 2.369 | Positive |

**Supplementary Table 2:** Effective Sample Size (ESS) for TransPhylo parameters.

| **Transmission cluster** | **Sampling proportion pi** | **Within-host coalescent rate Ne\*** | **Basic reproduction R** |
| --- | --- | --- | --- |
| **Cluster 1** | 1721 | 367 | 2407 |
| **Cluster 2** | 6027 | 1119 | 3400 |
| **Cluster 3** | 1890 | 391 | 3196 |
| **Cluster 4** | 4761 | 581 | 2649 |
| **Cluster 5** | 108783 | 933 | 10035 |
| **Cluster 6** | 2477 | 322 | 4910 |
| **Cluster 7** | 86600 | 568 | 3311 |
| **Cluster 8** | 3393 | 720 | 13012 |
| **Cluster 9** | 10640 | 202 | 17844 |
| **Cluster 10** | 2620 | 150 | 3533 |
| **Cluster 11** | 22204 | 1579 | 5094 |
| **Cluster 12** | 11576 | 8014 | 3128 |

**Supplementary Table 3:** Substitution rates for each transmission cluster

| **Transmission cluster** | **Median substitution rate (substitutions/site/year)** | **Median substitution rate (substitutions/genome/year)** |
| --- | --- | --- |
| **Cluster 1** | $4.3 \times 10^{-7}$ | 1.86 |
| **Cluster 2** | $2.1 \times 10^{-7}$ | 0.89 |
| **Cluster 3** | $1.4 \times 10^{-6}$ | 6.00 |
| **Cluster 4** | $8.8 \times 10^{-7}$ | 3.82 |
| **Cluster 5** | $3.5 \times 10^{-7}$ | 1.53 |
| **Cluster 6** | $9.1 \times 10^{-7}$ | 3.94 |
| **Cluster 7** | $1.2 \times 10^{-7}$ | 0.52 |
| **Cluster 8** | $2.0 \times 10^{-7}$ | 0.87 |
| **Cluster 9** | $3.2 \times 10^{-7}$ | 1.39 |
| **Cluster 10** | $2.4 \times 10^{-7}$ | 1.04 |
| **Cluster 11** | $1.5 \times 10^{-7}$ | 0.66 |
| **Cluster 12** | $1.2 \times 10^{-7}$ | 0.51 |

**Supplementary Figure 1:**
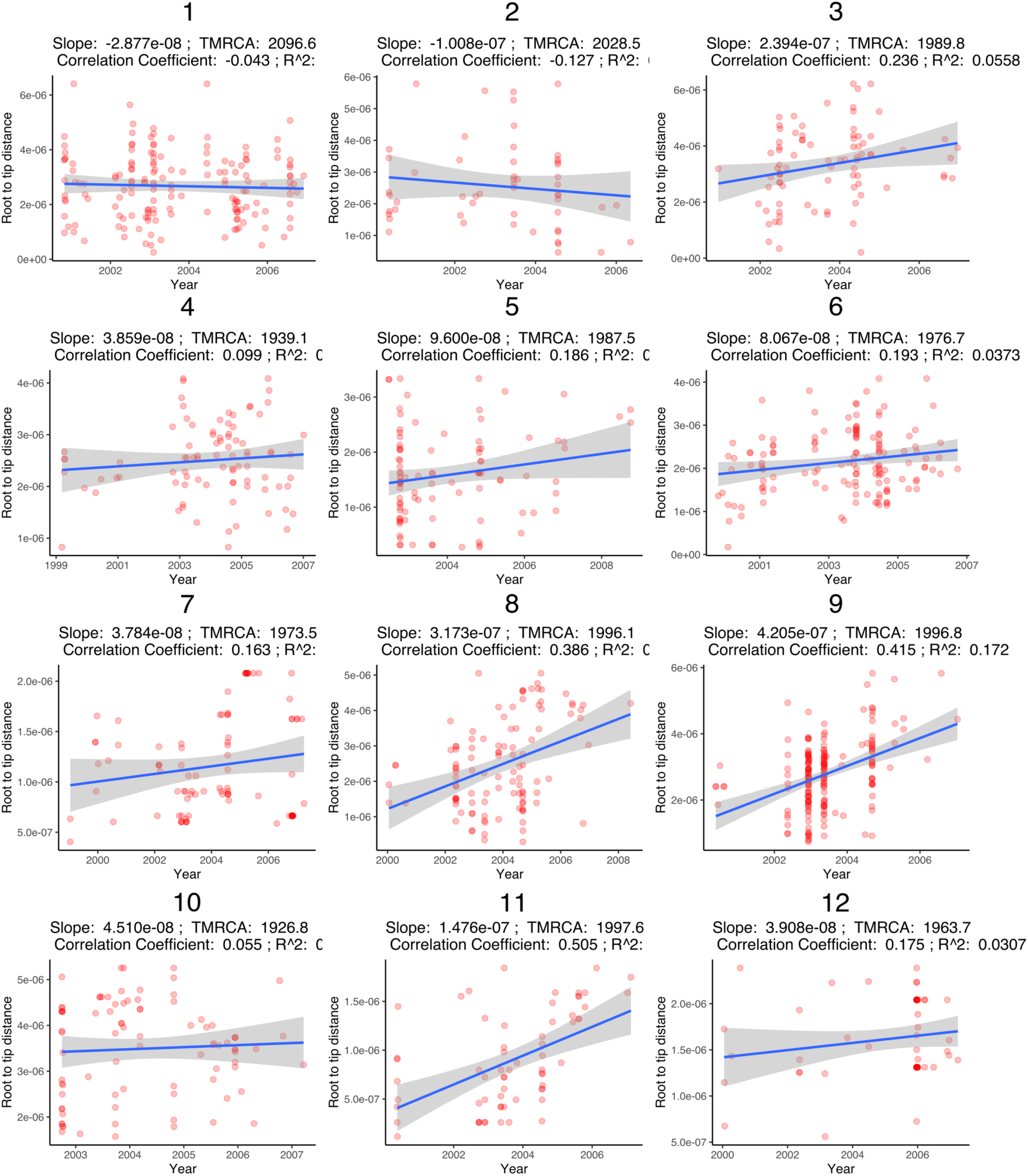
Root to tip distances plotted against sampling dates for all isolates in each transmission cluster.

**Supplementary Figure 2:**
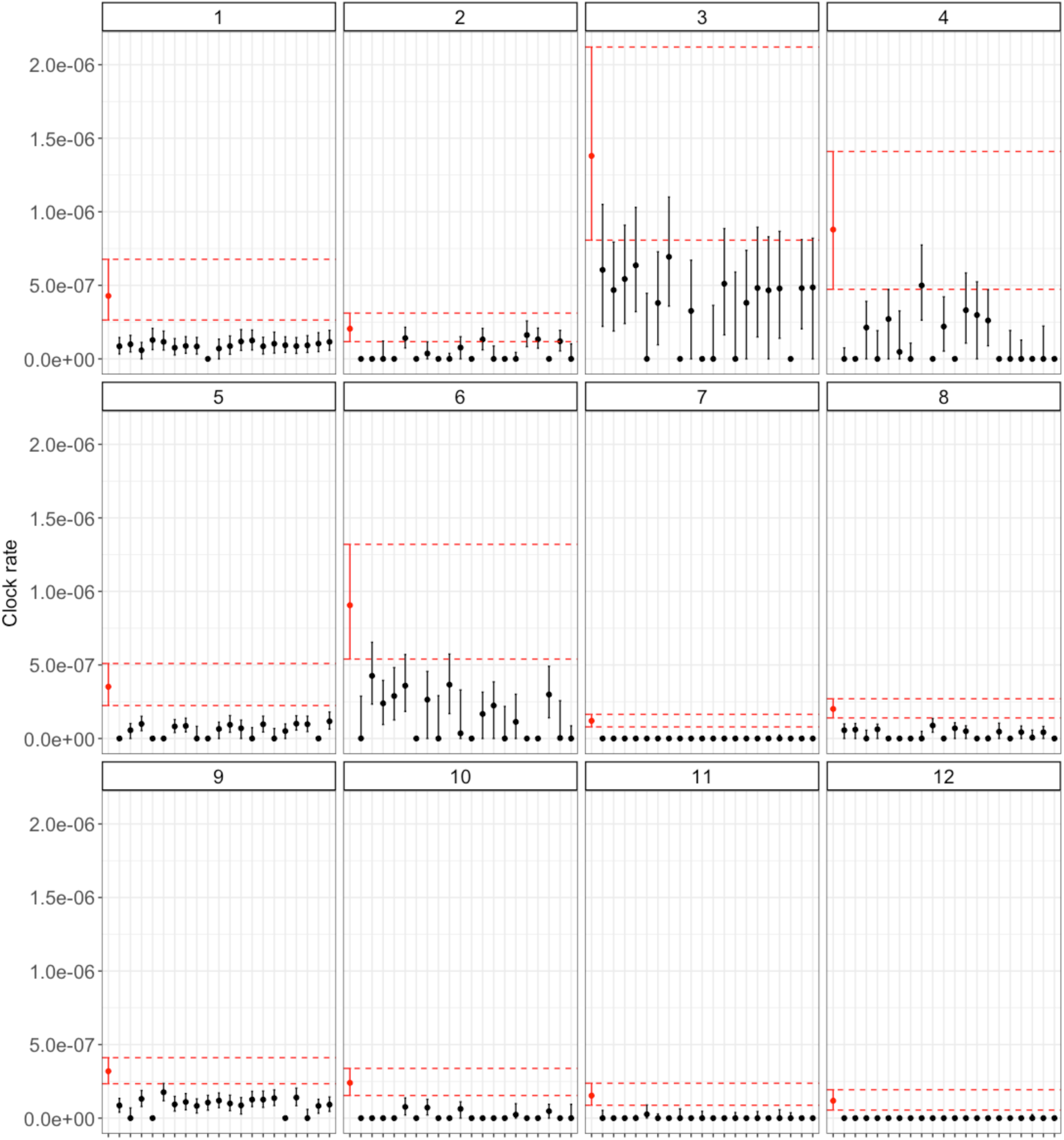
Date randomization (DTR) analysis in BEAST for each transmission cluster. Estimated substitution rates (mean and highest posterior density) shown in red for the observed dataset and black for the randomized datasets.

**Supplementary Figure 3:**
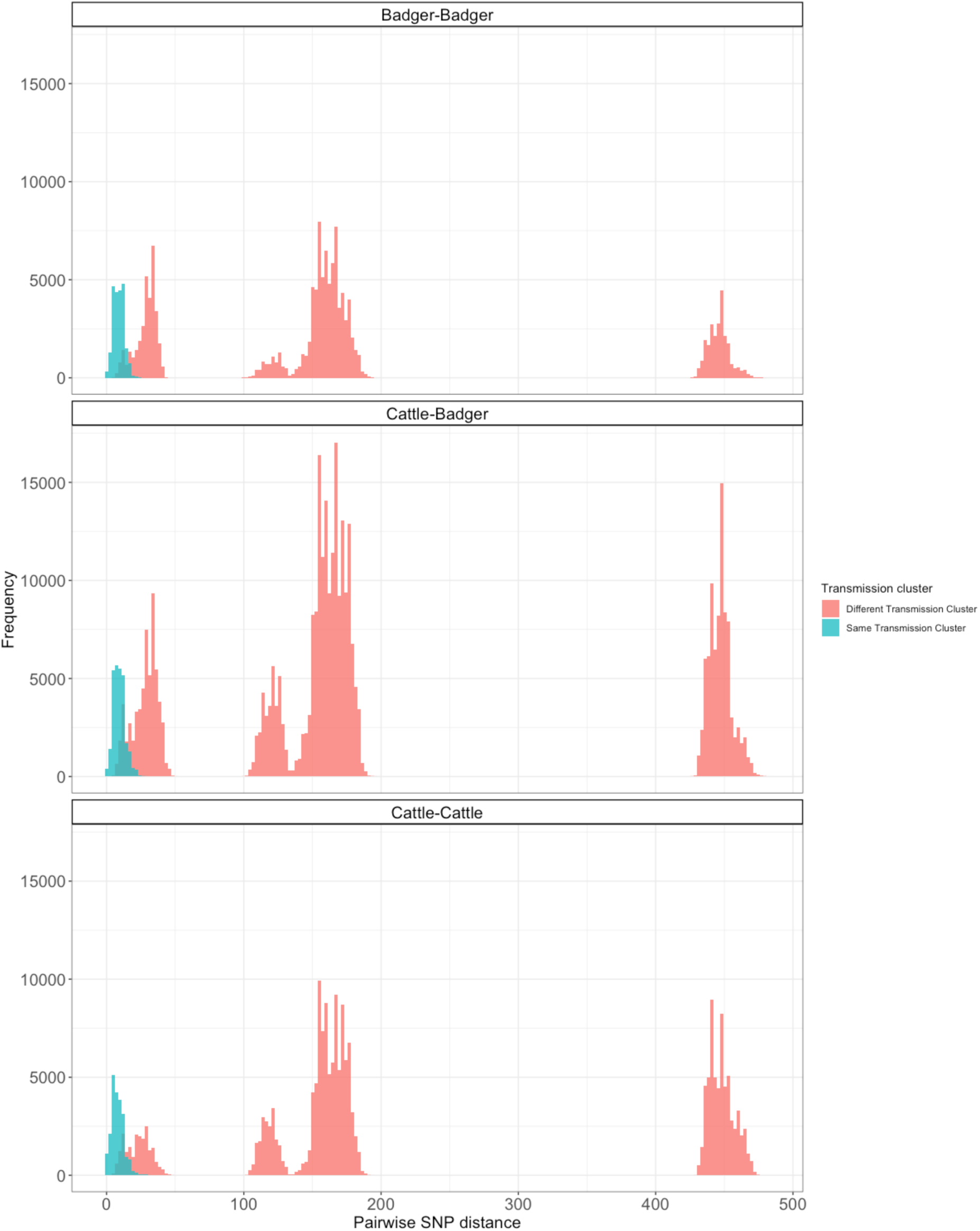
Pairwise distance histograms for all samples, coloured by between/within transmission cluster and separated by host pair.

**Supplementary Figure 4:**
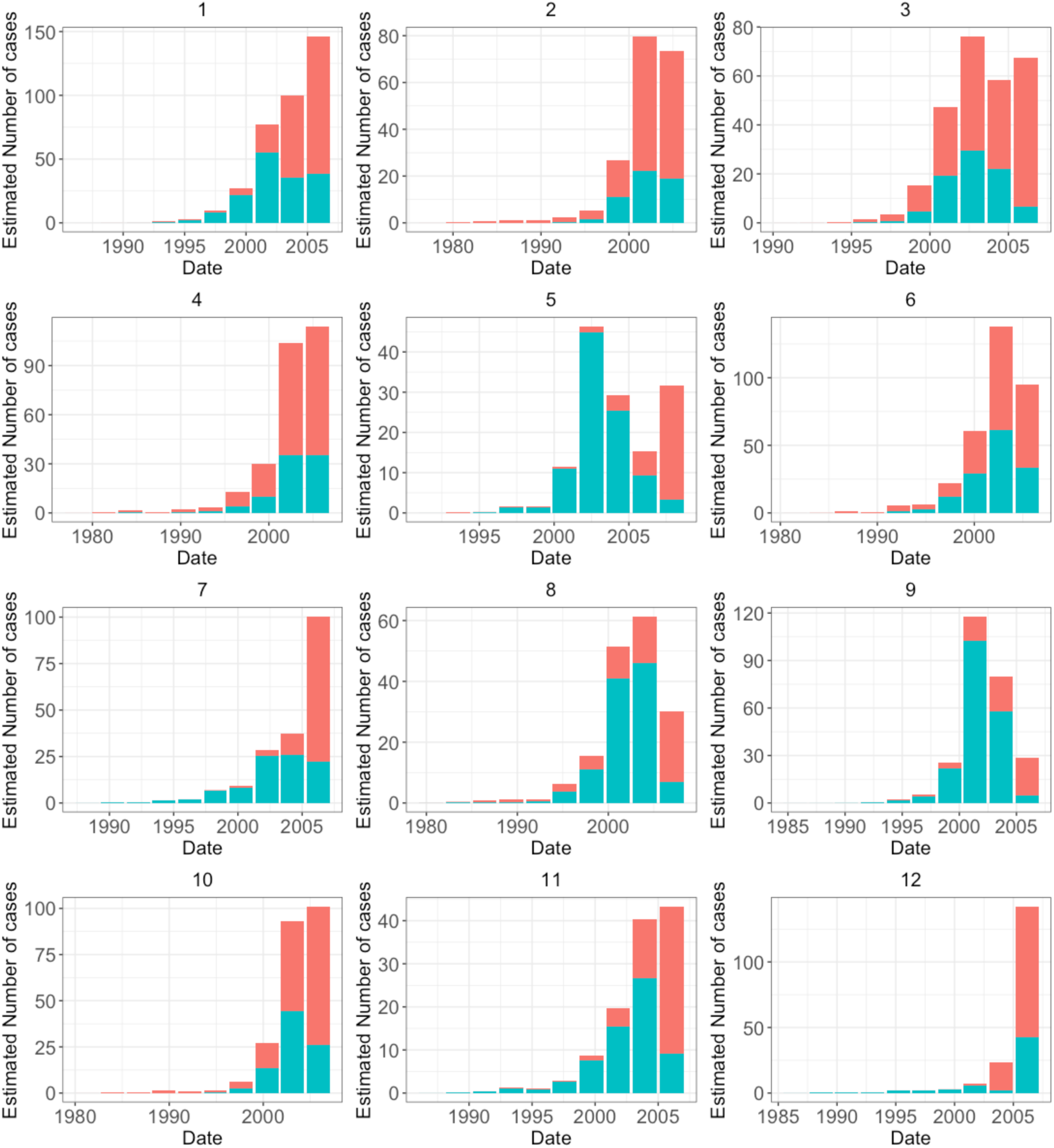
Proportion of sampled and estimated unsampled cases for each transmission cluster. Sampled and unsampled cases are shown in red and blue respectively

**Supplementary Figure 5:**
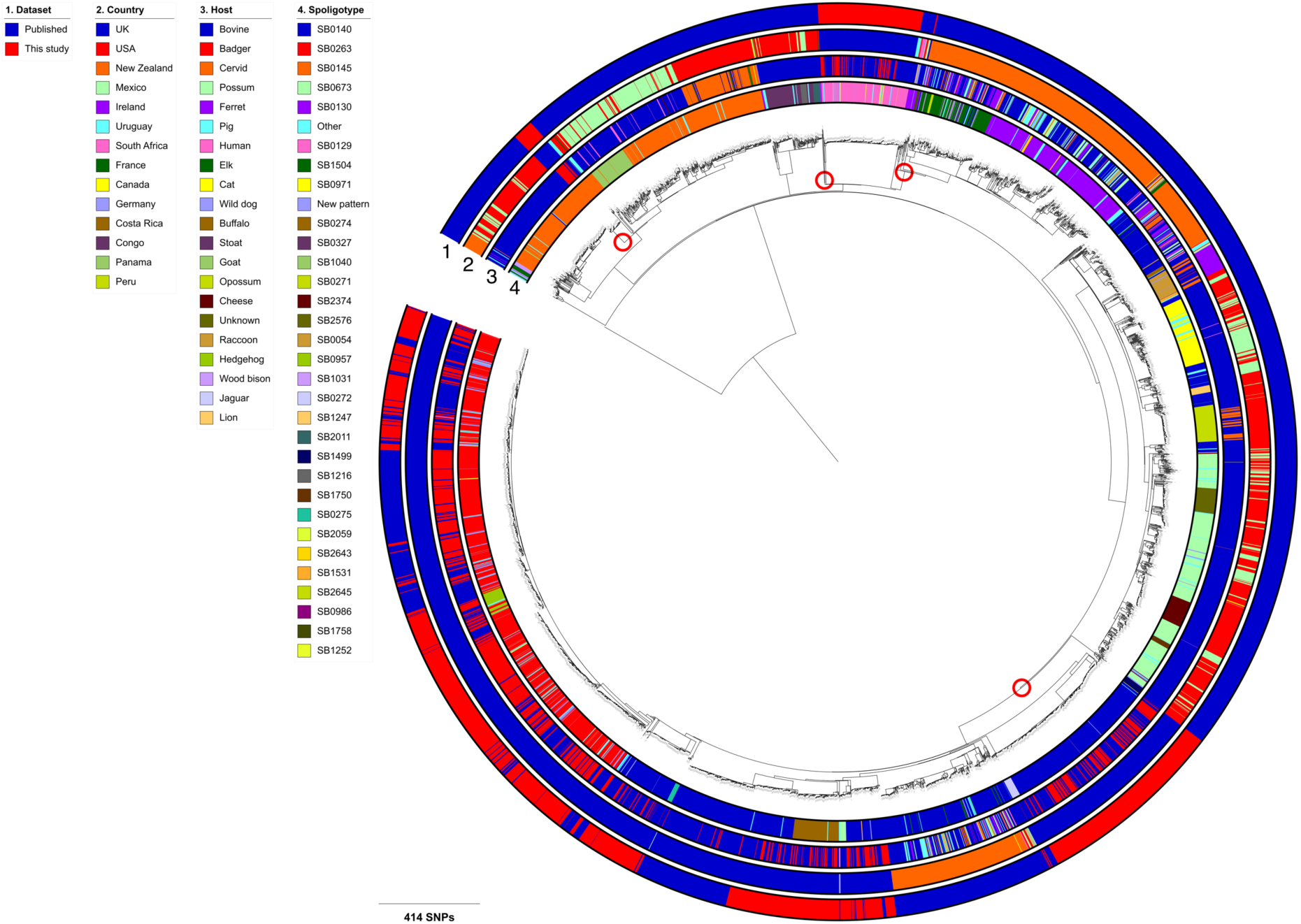
Maximum likelihood phylogenetic tree of 4,281 *Mycobacterium bovis* Eu1 isolates rooted with a *M. caprae* isolate as the outgroup. Dataset, country, host and spoligotype are shown as datastrips around the outside of the phylogenetic tree. Potential introductions of Eu1 into England are highlighted with red circles.

